# Neural Data Visualization for Scalable and Generalizable Single Cell Analysis

**DOI:** 10.1101/289223

**Authors:** Hyunghoon Cho, Bonnie Berger, Jian Peng

## Abstract

Single-cell RNA sequencing is becoming effective and accessible as emerging technologies push its scale to millions of cells and beyond. Visualizing the landscape of single cell expression has been a fundamental tool in single cell analysis. However, standard methods for visualization, such as t-stochastic neighbor embedding (t-SNE), not only lack scalability to data sets with millions of cells, but also are unable to generalize to new cells, an important ability for transferring knowledge across fast-accumulating data sets. We introduce net-SNE, which trains a neural network to learn a high quality visualization of single cells that newly generalizes to unseen data. While matching the visualization quality of t-SNE on 14 benchmark data sets of varying sizes, from hundreds to 1.3 million cells, net-SNE also effectively positions previously unseen cells, even when an entire subtype is missing from the initial data set or when the new cells are from a different sequencing experiment. Furthermore, given a “reference” visualization, net-SNE can vastly reduce the computational burden of visualizing millions of single cells from multiple days to just a few minutes of runtime. Our work provides a general framework for newly bootstrapping single cell analysis from existing data sets.

## Introduction

Complex biological systems arise from functionally diverse, heterogeneous populations of cells. Single-cell RNA sequencing (scRNA-seq) (Gawad et al., 2016), which profiles transcriptomes of individual cells rather than bulk samples, has been a key tool in dissecting the intercellular variation in a wide range of domains, including cancer biology (Y. Wang et al., 2014), immunology (Stubbington et al., 2017), and metagenomics (Yoon et al., 2011). scRNA-seq also enables the *de novo* identification of cell types with distinct expression patterns (Grün et al., 2015; Jaitin et al., 2014).

A standard analysis for scRNA-seq data is to visualize single cell gene expression patterns of samples in a low-dimensional (2D or 3D) space via methods like t-stochastic neighbor embedding (t-SNE) (Maaten and Hinton, 2008) or in earlier studies Principal Component Analysis (PCA) (Jackson, 2005), where each cell is represented as a dot and cells with similar expression profiles are located close to each other. Such visualization reveals the salient structure of the data in a form that is easy for researchers to grasp and further manipulate. For instance, researchers can quickly identify distinct subpopulations of cells through visual inspection of the image, or use the image as a common lens through which different aspects of the cells are compared. The latter is typically achieved by overlaying additional data on top of the visualization, such as known labels of the cells or the expression levels of a gene of interest (Zheng et al., 2017). While many of these approaches have initially been explored for visualizing bulk RNA-seq (Palmer et al., 2012; Simmons et al., 2015), methods that take into account the idiosyncrasies of scRNA-seq (e.g., dropout events where non-zero expression levels are missed as zero) have also been proposed (Pierson and Yau, 2015; B. Wang et al., 2017). Recently, more advanced approaches that visualize the cells while capturing important global structure such as cellular hierarchy or trajectory have been proposed (Anchang et al., 2016; Hutchison et al., 2017; Moon et al., 2017; Qiu et al., 2017), which constitute a valuable complementary approach to general-purpose methods like t-SNE.

Comprehensively characterizing the landscape of single cells requires a large number of cells to be sequenced. Fortunately, advances in automatic cell isolation and multiplex sequencing have led to an exponential growth in the number of cells sequenced for individual studies (Svensson et al., 2017) (Figure 1a). For example, 10x Genomics recently made publicly available a data set containing the expression profiles of 1.3 million brain cells from mice (10x Genomics, 2017). However, the emergence of such mega-scale data sets poses new computational challenges before they can be widely adopted. Many of the existing computational methods for analyzing scRNA-seq data require prohibitive runtimes or computational resources (10x Genomics, 2017). In particular, the state-of-the-art implementation of t-SNE (Van Der Maaten, 2014) requires 1.5 days to run on 1.3 million cells based on our estimates.

**Figure 1:**
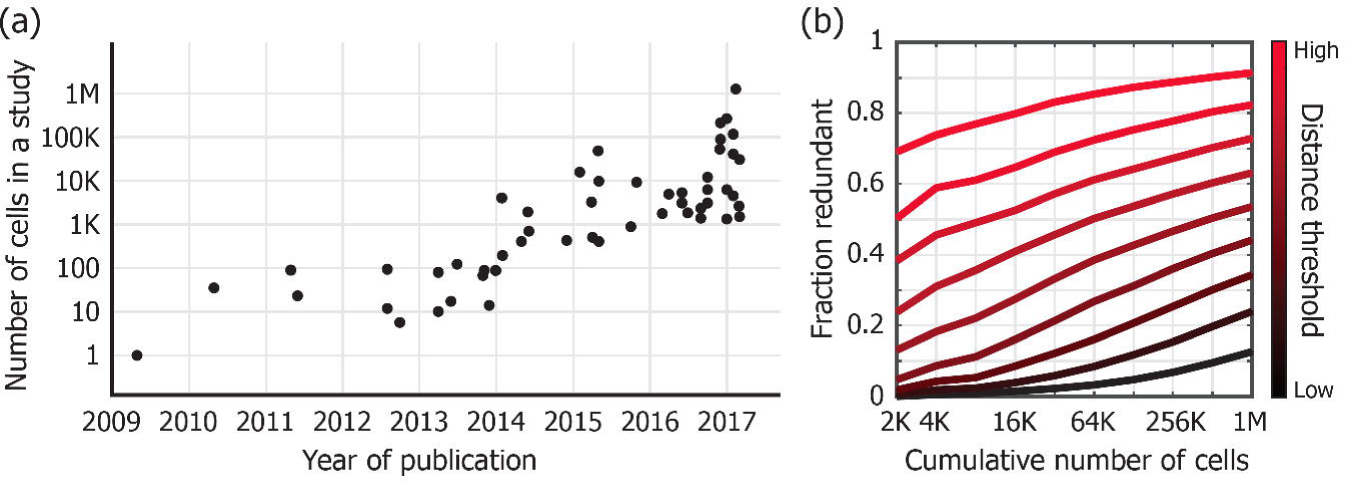
The increasing scale and redundancy of single-cell RNA-seq data sets. (a) The exponential increase in the number of single cells sequenced by individual studies (adapted from Svensson et al. (2017) with a new point for the 1.3 million cell data **(10x Genomics, 2017)**. Note that the y-axis scales exponentially. (b) Retrospective analysis of redundancy in the 1.3 million cell data **(10x Genomics, 2017)** with 2000 initial cells and repeated doubling of the data size. For each batch added, we computed the distribution of the cells’ minimum Euclidean distance to cells already observed based on their gene expression. Each curve corresponds to a particular distance threshold for deeming the new cell redundant. The thresholds are chosen as the deciles of the overall distribution of minimum Euclidean distances.

Here, we introduce *neural t-SNE* (net-SNE), a scalable and generalizable method for visualizing single cells for scRNA-seq analysis. Taking inspiration from compressive genomics (Loh et al., 2012), we exploit the intuition that when a large number of cells are sequenced, a significant portion of the cells are redundant (i.e., highly similar to other cells). Taking advantage of the expressive power of neural networks (NNs), which has been demonstrated in numerous applications (LeCun et al., 2015), net-SNE trains a NN to learn a high-quality mapping function that takes an expression profile as input and outputs a low-dimensional embedding in 2D or 3D for visualization. Unlike t-SNE, the mapping function learned by net-SNE can be used to map previously unseen cells that were not included in the input data. This capability allows for novel workflows for single cell studies, where newly observed cells are visualized in the context of existing data sets to gain additional insights.

To demonstrate visualization quality as well as scalability, we show that net-SNE learns visualizations that are similar to those of t-SNE on 14 scRNA-seq data sets of various cell types and data sizes up to 1.3 million cells. Next we focused on generalizability, demonstrating that net-SNE newly achieves the ability to map previously unseen cells; in particular, we show that net-SNE can not only identify subtypes of cells that were not included in the initial data, but also be used to bootstrap the visualization from a subset of data to achieve significantly better scalability to mega-scale data sets. Given the inherent redundancy in biological data (Yu et al., 2015), we expect our techniques for neural data visualization to accelerate and enhance other high dimensional biological data analyses beyond visualization.

## Results

### Increasing redundancy in single cell data sets

We first set out to empirically assess the extent to which additional sequencing of single cells from the same biological source capture unforeseen expression patterns. Starting with 2,000 randomly chosen cells from the 10x Genomics scRNA-seq data set with 1.3 million mouse neurons (Brain1m), we repeatedly doubled the data size up to a million cells by sampling the remaining cells (without replacement) and measured how redundant the newly added cells are compared to the ones already observed (Figure 1b). As the scale of data grows, sequencing more cells exhibits a clear diminishing return in terms of capturing cells with unique expression patterns. For example, the majority of cells (53%) in the final half of the data can be considered redundant according to a certain distance threshold, which deems only 10% of the cells redundant in the initial batch. Nevertheless, a considerable fraction of the newly-observed cells remains unique even at the scale of a million cells. For instance, 10% of the cells in the final half are as unique as the top 30% of the initial batch, which suggests that the push toward a higher cell count is indeed valuable for gaining access to relatively unexplored regions of the gene expression landscape, albeit with decreasing effectiveness.

These results imply that, as researchers collectively accumulate scRNA-seq data for a particular biological system (e.g., tissue, organism, or microbial population) to the scale of millions of cells, a significant portion of newly sequenced cells will fall into the space already visited by existing data, where useful insights may be available from previous analyses. This observation motivates our development of net-SNE, which allows new data to be mapped onto an existing visualization to accelerate such knowledge transfer across different studies or experiments.

### Overview of net-SNE

net-SNE achieves generalizability by training a neural network to learn a parameterized embedding function that takes a cell’s expression profile as input and outputs the coordinates in a low-dimensional space for visualization (Methods). Given the wide success of t-SNE in single cell biology (Amir et al., 2013), we aim to emulate the behavior of t-SNE while newly achieving the ability to map new cells, by training our neural network to optimize the same objective function as t-SNE. This objective function intuitively captures how faithfully the local structure among the input vectors (i.e., single cell gene expression profiles) are represented in the visualization. Although our *parametric* approach to t-SNE has been theoretically considered (Van Der Maaten, 2009), its application to real-world, large-scale data sets have been considerably limited due to the difficulties in successfully training a neural network for a complex task like t-SNE. Our work employs new optimization techniques to improve the scalability of neural network training for t-SNE (Methods) and also newly demonstrates the effectiveness of this approach for enabling large-scale and translational single cell analysis.

### net-SNE learns high quality visualizations of single cells

To evaluate the ability of net-SNE to accurately model the visualization of single cell data sets, we tested it on 13 existing scRNA-seq data sets of varying sizes *with known clusters* (Methods). We found that for all of the data sets net-SNE is able to learn an embedding that closely matches the output of t-SNE (Figures 2a, S1).

**Figure 2:**
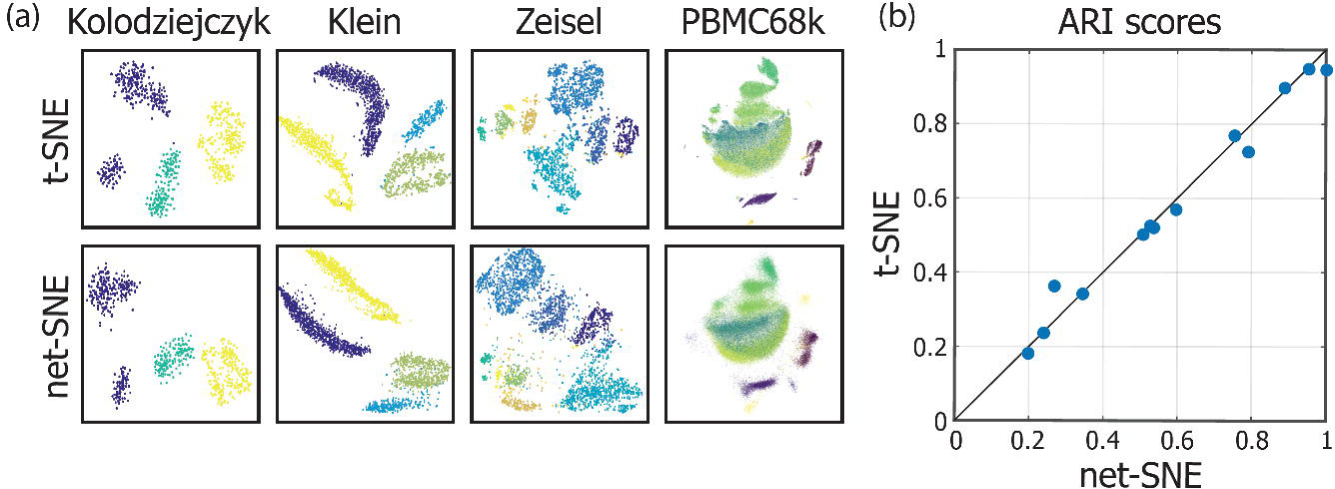
net-SNE recapitulates t-SNE mapping on 13 benchmark data sets with known subtypes. (a) Comparison of net-SNE and t-SNE visualizations on four largest benchmark data sets with known clusters. Colors indicate known cell types provided by the original work. Figures for the remaining data sets are provided in Figure S1. (b) Quality of each visualization is quantified by the adjusted Rand index (ARI) between the known labels and the output of k-means clustering based on the embedding. Each dot represents one of the 13 data sets analyzed. Results based on agglomerative clustering instead of k-means are provided in Figure S2, also showing a high concordance between net-SNE and t-SNE.

To systematically evaluate the quality of embeddings produced by net-SNE, we assessed the agreement between the known subtypes and sought to computationally identify clusters based on the low-dimensional embeddings. The level of agreement was quantified by the adjusted Rand index (Rand, 1971), following previous work (Kiselev et al., 2017). We obtained the clusters by applying the standard k-means clustering algorithm (Hartigan and Wong, 1979) to the embeddings, where the known number of clusters was provided to the algorithm. As shown in Figure 2b, net-SNE achieves clustering accuracy that is comparable to t-SNE for all 13 data sets, which agrees with the visual concordance of the two methods. An analogous analysis we performed, based on agglomerative clustering instead of k-means, leads to similar concordance between net-SNE and t-SNE (Figure S2).

Notably, we obtained all of these results using a relatively simple neural network with only two layers of nonlinearities with 50 units each. In additional experiments, not only do we observed that the net-SNE results are reasonably stable across a wide range of these parameters, we also found that the size of our network can be reduced to as low as 10 units per layer without significantly sacrificing the quality of visualizations (Figure S4), even for the PBMC68k data set containing tens of thousands of cells. This finding suggests the relationship between gene expression profiles and the clustering pattern of cells is simple enough to admit a concise characterization across a wide range of data sets.

### net-SNE accurately maps new cells

To demonstrate the potential of net-SNE for translational analyses, we performed a cross validation experiment where an entire cluster of cells was removed from a data set and placed onto the visualization after the fact. While the original t-SNE does not support the visualization of new data points, we considered as baseline a naïve extension of t-SNE where the embedding of a new cell is determined as the average position (in the low-dimensional space) of the cell’s five nearest neighbors in the initial data according to expression measurements (t-SNE+k-NN). An alternative extension of t-SNE where the new cells are randomly initialized and optimized while fixing the positions of the initial cells, similarly lacks the scalability to mega-scale datasets as the original t-SNE and thus was not considered in our analysis.

Figure 3 shows our cross validation results on the Klein data set (Klein et al., 2015), which contains four known clusters, each of which was held out in four separate experiments. Remarkably, in three out of the four cases, the embedding learned by net-SNE accurately positioned the held-out cells as a distinct cluster, despite the fact that the training data did not contain any of the cells from this cluster. In contrast, our nearest neighbor-extension of t-SNE (t-SNE+k-NN) overlaid most of the new cells onto existing clusters and ended up incorrectly outputting an obfuscated map. Although visualizing the entire data set from scratch results in higher quality scores than both approaches for generalization (Figure S4a), we note that the initial generalization can be further optimized by net-SNE or t-SNE to correct for the noise introduced during the generalization.

**Figure 3:**
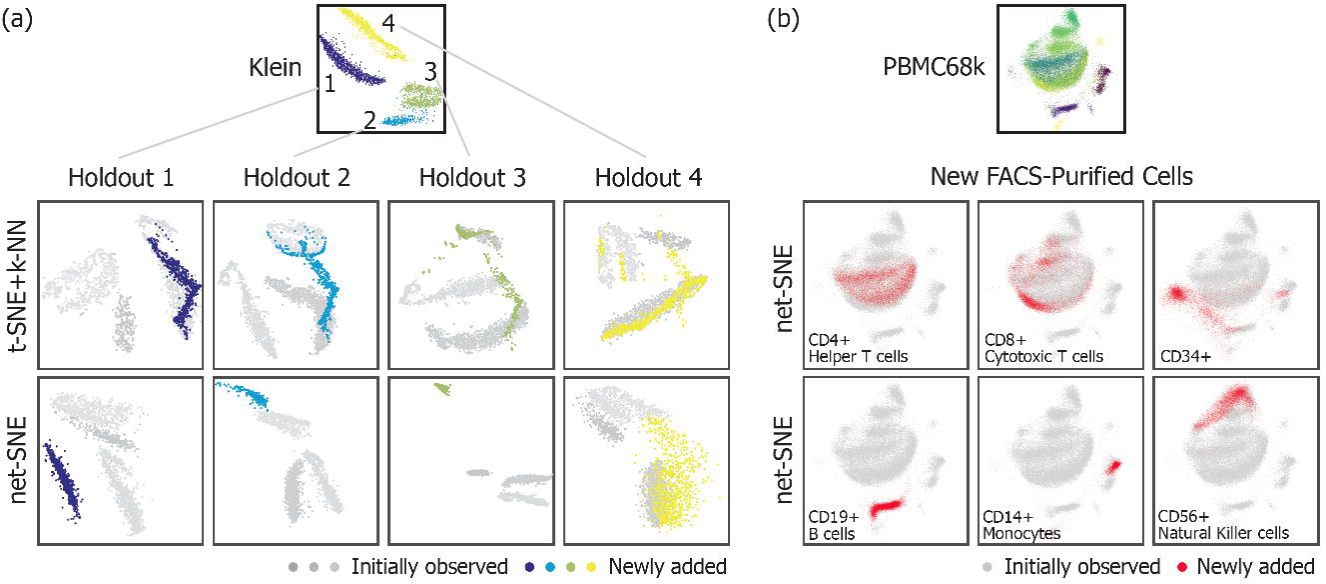
net-SNE generalizes to unseen cells. (a) Each column represents a cross validation experiment where the visualization is performed only on a subset of data (shown in grey), where all cells corresponding to one of the four known cell types in the Klein data set were held out. Held-out cells were later added to the visualization (shown in color). We compared net-SNE to a naïve extension of t-SNE where each new cell is placed at the average position of the five nearest neighbors in the initial data set. Unlike t-SNE, net-SNE is able to identify the newly added cells as a separate cluster. Also, in the setting where the new cells belong to subtypes already represented in the initial data set, net-SNE accurately assigns the new cells to the respective clusters (Figure S4b). (b) To further test net-SNE’s generalizability, we obtained scRNA-seq data sets from 10x Genomics for six representative blood cell types (shown in subfigure), each purified via fluorescence activated cell sorting (FACS). We visualized each of the purified cell populations (in red) over a pre-trained net-SNE embedding of the PBMC68k data set (shown in gray), which is from a whole blood sample encompassing all of the purified cell types. Despite coming from a different sequencing experiment and different sample preparation, many of the purified cell types were precisely mapped by net-SNE to well-defined clusters in the visualization, demonstrating its potential for cross-data-set knowledge transfer.

The setting of this cross validation analysis may arise in practice where a rare subpopulation of cells was omitted from the initial data set due to small data size. Our results suggest that, while the naïve nearest neighbor-based projection of newly observed cells (including the rare subtype) will likely render the new subtype invisible, net-SNE is still able to identify the new cluster, given that its gene expression is sufficiently distinct from that of existing cells. Also we confirmed that, when the new cells are from a subtype that is already represented in the initial data set, net-SNE is able to accurately assign the new cells to the correct cluster (Figure S4b).

To further demonstrate the utility of net-SNE’s generalization performance beyond cross validation, we obtained scRNA-seq data sets of six different purified blood cell types (CD4+, CD8+, CD14+, CD19+, CD34+, and CD56+) from 10x Genomics (Methods) and projected each data set onto a pre-trained, net-SNE visualization of the PBMC68k data set, which consists of cells obtained from a whole blood sample in which all of the aforementioned cell types are present. Despite the differences in sample preparation and the possibility of batch effects (Tung et al., 2017), net-SNE accurately positions the new cells onto existing clusters, which immediately reveals meaningful properties of a number of clusters in the PBMC68k data set (Figure 3b). We further confirmed that the information we elucidated using this approach is consistent with the characterization provided in the original publication of PBMC68k data (Zheng et al., 2017).

### net-SNE accelerates visualization of millions of cells

After validating net-SNE’s ability to generalize to new cells, we then asked whether this ability can be exploited to achieve fast visualization of mega-scale data sets. Drawing from the intuition that data sets of this scale can be accurately represented by a smaller subset of cells due to high redundancy, we first trained net-SNE on a subset of 100K cells from the Brain1m data set containing 1.3 million cells and later applied the learned embedding function to the entire data set. This fast approach took around only 20 minutes overall and resulted in a higher quality map than the output of t-SNE with the default parameter settings, which took 13 hours to finish. Note that we use the Kullback-Leibler (KL) divergence objective score—the quantity minimized by both t-SNE and net-SNE—as the metric of quality (inversely related), which is more objective than a visual assessment. If a researcher already has access to a pre-trained mapping based on an existing data set, then the reduction in runtime achieved by net-SNE is likely to be even more drastic (e.g., days for t-SNE to a few minutes).

It is worth noting that the original data set provided by 10x Genomics also included a t-SNE embedding, which appeared higher in quality than the t-SNE output we obtained using the default setting (Figure 4e). While the objective score we computed based on the published embedding was superior to net-SNE’s initial generalization, 45 minutes of further optimization of net-SNE was sufficient to outperform this score (Figure 4c).

**Figure 4:**
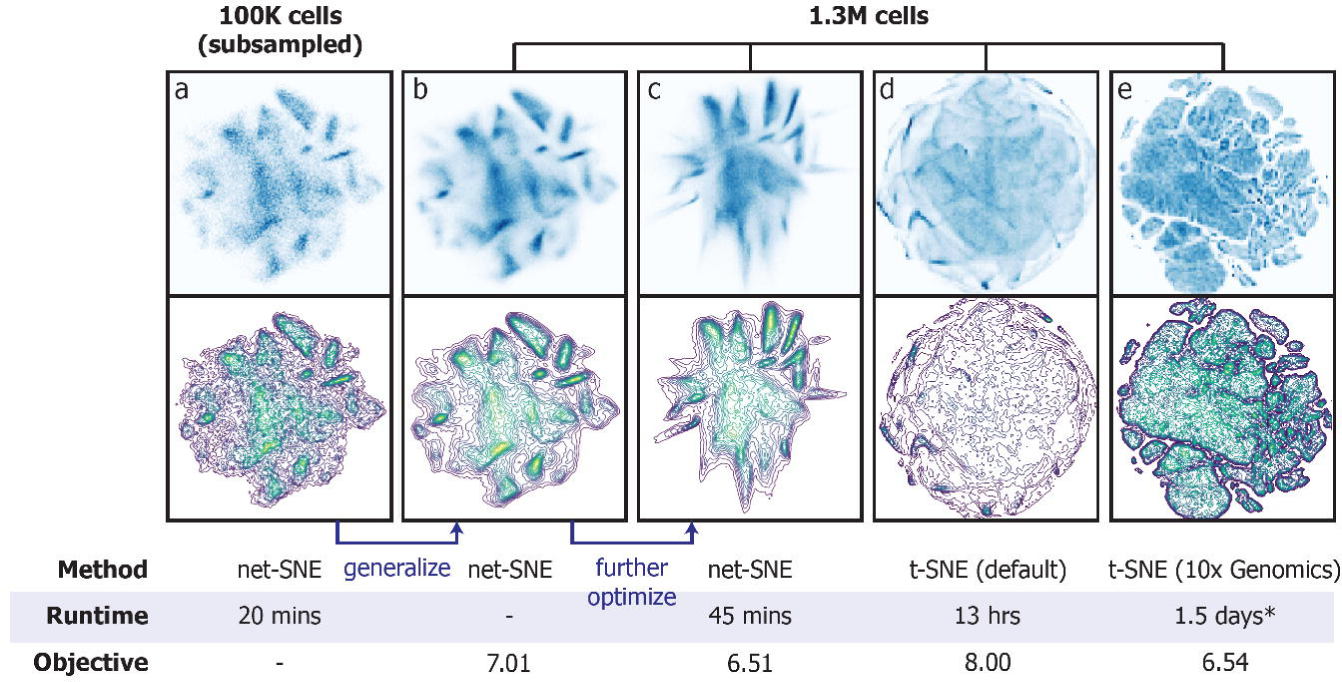
net-SNE enables fast visualization of mega-scale data sets. Brain1m data set with 1.3 million cells is visualized using a novel bootstrap approach enabled by net-SNE, where the embedding learned on a subset of 100K cells (a) was used to initialize training on the whole data set. Our initial generalization to the full data instantly obtained (in minutes) a visualization (b) of higher quality, as measured by the KL divergence objective score minimized by both methods (Methods), than that of t-SNE with default parameters achieved after 13 hours (d). While the t-SNE embedding provided in the original data set by 10x Genomics (e) achieves a better objective, net-SNE outperforms this embedding with less than an hour of further training (c). Top row shows the heat map for each embedding using a linear color map, where the highest value represented is chosen for each plot to achieve the best clarity. Bottom row shows contour plots of the same data. Lower objective score corresponds to better agreement between the gene expression landscape and the visualization. *The visualization by 10x Genomics can be closely reproduced by increasing the number of iterations for t-SNE, and based on our experiments t-SNE required 1.5 days to achieve a solution with a comparable score.

The visual difference between the net-SNE and t-SNE outputs in this experiment can be partially attributed to the fact that there are likely many locally optimal solutions to the same optimization problem solved by both methods. Since net-SNE was initialized with a specific solution as opposed to the random initialization of t-SNE, it is understandable that they converge to different solutions. Although the clusters appear to be more clearly separated in the t-SNE output, the fact that our visualization actually achieves a better objective score suggests the possibility of the abrupt boundaries in t-SNE being a potential artifact that is not warranted by the underlying gene expression values. Since net-SNE restricts the space of possible visualizations to those that can be modeled as a continuous function as specified by our relatively simple neural network, it has a tendency to obtain a “smoother” visualization, which could potentially be a more accurate representation of the underlying data than t-SNE output given the continuity of gene expression levels. Note that we obtained our results on the Brain1m data set using a two-layer neural network with 50 hidden units per layer, as in our aforementioned analyses.

After observing that we can achieve a visualization similar to the one provided by 10x Genomics by increasing the number of iterations for t-SNE (thus increasing runtime), we performed an experiment where we ran t-SNE until it reaches an objective score matching the provided embedding. This resulted in a runtime estimate of 1.5 days for the published visualization, which is substantially longer than that required by net-SNE to achieve a superior performance (i.e., 20 minutes of pre-training and 45 minutes of further optimization). Notably, if the generalization by net-SNE was performed based on a well-characterized data set, then the map obtained by net-SNE would have the additional benefit of allowing researchers to immediately transfer insights from the existing data.

## Discussion

As we enter the age of mega-scale single cell analysis, new computational methods that take advantage of the growing redundancy in the scRNA-seq data are needed. To this end, we have presented a new visualization method, net-SNE, which uses a neural network to learn a parametric embedding function that emulates t-SNE’s visualization while newly achieving the ability to map previously unseen cells. We have demonstrated that net-SNE not only learns high quality maps like t-SNE, but also gracefully generalizes to unseen cells—even when a whole subpopulation is missing from the initial data set or when the new data comes from a different biological sample. Indeed, net-SNE’s ability to generalize allows researchers to exploit redundancy across different data sets by projecting the cells in one data set onto another to facilitate transfer of knowledge. In addition, we have shown that using a pre-trained embedding from a subsampled (or an existing) data set is an effective way for performing fast visualization of mega-scale scRNA-seq data sets. Our approach achieves significantly better scalability than t-SNE, which is on the verge of being impractical for data sets with more than a million cells. Although a number of recent studies introduce new techniques for improving the scalability of data visualization tools (Dzwinel and Wcisło, 2015; Tang et al., 2016), they do not address the lack of generalizability that net-SNE overcomes. Notably, many single cell analysis methods that are newly being developed can be used in conjunction with net-SNE to potentially achieve enhanced visualization quality; for instance, methods that account for dropout events (B. Wang et al., 2017) or batch effects (Haghverdi et al., 2017) can be performed to obtain an improved input similarity matrix before net-SNE is applied.

The fact that net-SNE obtains a visualization where the coordinates are directly modeled by the neural network as a function of gene expression opens up new directions for further research. In particular, one can investigate the parameters of the neural networks trained by net-SNE for insights into what types of expression patterns are being utilized for t-SNE-like visualizations. Furthermore, while the characterization of conspicuous clusters in t-SNE output has typically been done by summarizing the expression of cells that belong to the cluster of interest, net-SNE enables a more direct and potentially more effective approach that analyzes the behavior of the embedding function.

A recently proposed idea of building a reference map of all human cell types based on high-throughput single-cell experiments (called the Human Cell Atlas (Regev et al., 2017)) is closely related to our vision. While embedding all cell types in a space with as few as two or three dimensions is unlikely to be successful given the complexity of the problem, we believe that insights from net-SNE and its future extensions may lead to an effective approach for learning reduced vector space representations of all human cells that can be readily used with new and existing computational methods to further advance our understanding of biology.

## Author Contributions

Conceptualization, H.C., B.B., and J.P.; Data curation, H.C. and J.P.; Investigation, H.C. B.B., and J.P.; Methodology, H.C., B.B., and J.P.; Resources, B.B.; Software, H.C.; Validation, H.C.; Visualization, H.C.; Writing – Original Draft, H.C.; Writing – Review & Editing, H.C., B.B., and J.P.; Funding acquisition, B.B.; Supervision, B.B. and J.P.

## Acknowledgements

A single page abstract of this work will appear in RECOMB 2018. This work was partially supported by NIH grant R01GM081871.

## Methods

### Review of t-Stochastic Neighbor Embedding

Let ***x***_1_, …, ***x***_*n*_ ∈ ℝ^*d*^ represent the (normalized) expression profiles for each of the *n* cells in a scRNA-seq data set that we wish to visualize in ℝ^*s*^, where *d* is typically on the order of tens of thousands (number of human genes) and *s* is two or three. More precisely, we want to learn the low-dimensional embedding of the cells ***y***_1_, …, ***y***_*n*_ ∈ ℝ^*s*^ that capture the low-dimensional structure represented by the original input vectors ***x***_1_, …, ***x***_*n*_.

A widely-used approach called t-stochastic neighbor embedding (t-SNE) (Maaten and Hinton, 2008) relates the notion of *quality* of an embedding ***y***_1_, …, ***y***_*n*_ (inversely) to the Kullback-Leibler (KL) divergence between the two probability distributions ***P*** and ***Q*** defined over all pairs of cells, which reflect how the cells are laid out in the input and output (embedding) spaces, respectively. The probability assigned to a particular pair (*i*, *j*) in each distribution represents how close the two associated vectors are---i.e., (***x***_*i*_, ***x***_*j*_) for ***P*** and (***y***_*i*_, ***y***_*j*_) for ***Q***. Intuitively, maximizing the agreement between ***P*** and ***Q*** corresponds to finding a good embedding that faithfully represents the structure in the original data.

Formally, t-SNE solves the following optimization problem

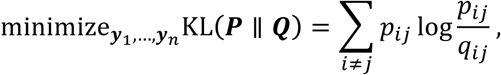

where

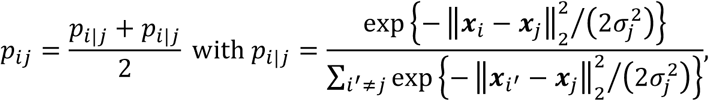

and

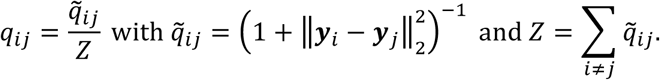

Note *p*_*ij*_ and *q*_*ij*_denote the (*i*, *j*) element of matrices ***P*** and ***Q***, respectively. In addition, *σ*_*j*_ is a parameter that is tuned for each *j* to ensure *p*_*i|j*_ achieves a predefined value of information-theoretic entropy.

t-SNE solves the above optimization problem via gradient descent on the embedding vectors ***y***_1_, …, ***y***_*n*_ with random initialization. As derived in the original paper (Maaten and Hinton, 2008), the gradient for each ***y***_*i*_ is given as

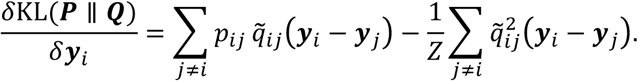

Computing this gradient for every cell *i* would require *O*(*n*^*2*^) computation, which is prohibitive for large *n*. In the state-of-the-art implementation of t-SNE (Van Der Maaten, 2014), this expression is approximated in two ways for computational efficiency. First, ***P*** is approximated with a sparse matrix based on k-nearest neighbors for each cell, which greatly speeds up the computation of the first term since most summands are zero. Second, an efficient data structure (space-partitioning trees (Samet, 1984)) is built over ***y***_1_, …, ***y***_*n*_ so that the second summation can be coarsely approximated by grouping terms corresponding to nearby ***y***_*i*_’s together. Even with these optimizations, applying t-SNE to data sets with millions of cells requires days of computation as shown in our results.

### Our Method: Neural t-SNE

We introduce neural t-SNE (net-SNE), which models each embedding vector ***y***_*i*_ as the output of a parameterized embedding function evaluated at the corresponding input vector ***x***_*i*_. Importantly, our approach is generalizable—i.e., it induces the embedding of any point in the input space, not just the observed data points as in t-SNE. We use standard feedforward neural networks (NNs) (LeCun et al., 2015) to represent the embedding function, drawing from the intuition that NNs have sufficient expressive capacity to find high-quality maps similar to those typically uncovered by t-SNE.

The precise form of the parameterized mapping of net-SNE is as follows. Let *ℓ* be the number of hidden layers in the NN and *u* be the number of units in each layer (same for every layer). Furthermore, let ***W***^(*t*)^ ∈ ℝ^*u×u*^ (ℝ^*u×d*^ for *t* = 1) be the weight matrix and ***b***^(*t*)^ ∈ ℝ^*u*^ be the intercept associated with layers *t* = 1, …, *ℓ*. An additional weight matrix ***W***^*(ℓ+1)*^ is associated with the final output layer. Given a data point ***x***_*i*_, the forward pass through the NN to compute the embedding ***y***_*i*_ can be recursively described as

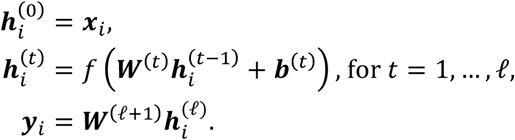

Note that *f* denotes an element-wise nonlinear activation function (e.g., sigmoid or rectifier). In the following, we compactly represent the above NN-based embedding function as ***y***_*i*_ = NN(***x***_*i*_; *Θ*), where *Θ* refers to the network parameters ***W***^*(1)*^, …, ***W***^*(ℓ+1)*^ and ***b***^*(1)*^, …, ***b***^*(ℓ)*^.

Given a NN that defines an embedding for every point in the input space, net-SNE optimizes the same KL divergence objective as t-SNE over the observed data points, via gradient descent. To see how the gradients are computed in net-SNE, first note that for a particular network parameter *θ* ∈ *Θ*, we have

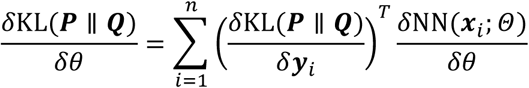

by the chain rule and using ***y***_*i*_ = NN(***x***_*i*_; *Θ*). Notice the first term in each product is identical to the t-SNE gradient and can be computed in the same manner. The second term can be computed via standard backpropagation algorithm (LeCun et al., 2015).

Intuitively, here we keep most of the computation in t-SNE intact, but add an additional step to each iteration, where, after computing the gradients for ***y***_1_, …, ***y***_*n*_ as in t-SNE, we propagate them backward through the NN to update the network weights accordingly. Consequently, net-SNE is compatible with any computational optimization for t-SNE. In particular, our implementation of net-SNE incorporates the state-of-the-art version of t-SNE based on the Barnes-Hut approximation (Van Der Maaten, 2014) to the gradients with respect to ***y***_1_, …, ***y***_*n*_, which achieves substantially faster runtime than vanilla t-SNE.

Although our method was independently developed, we note that a theoretical approach of training a parametric embedding (e.g., a neural network) for t-SNE via the chain rule has been previously described in an earlier work (Van Der Maaten, 2009). However, given the difficulty in successfully training a neural network to find good solutions to the t-SNE objective on large-scale data sets, practical adoption of this approach has been limited. In our work, we introduce additional techniques described in the following section to improve the effectiveness of neural network-based visualization, while also demonstrating its utility for single cell analysis on a wide range of benchmark data sets.

### Accelerating net-SNE via Stochastic Optimization

To fully exploit the generalizability of net-SNE, we improve upon the above procedure with techniques from stochastic optimization. First, note that the gradient for *θ* ∈ *Θ* given in the previous section is a summation over all data points, and thus can be approximated with a randomly chosen subset ℬ ⊂ {1, …, *n*} (“mini-batch”) as

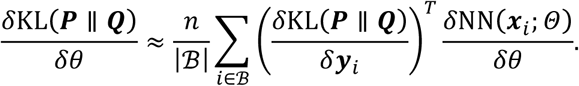

Similarly, the gradient with respect to ***y***_*i*_ for *i* ∈ ℬ can be approximated as

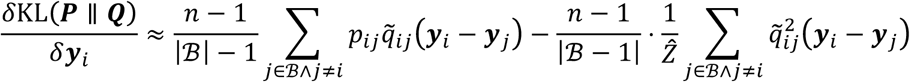

using the approximate normalization factor

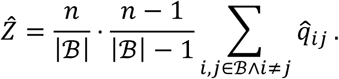

With small |ℬ| and precomputed ***P*** (which typically constitutes a small fraction of the runtime of t-SNE), this approach greatly reduces the required computation in each gradient step and improves the rate of convergence to a good visualization, as we demonstrate in our experiments. Our observation is in line with well-known results in the field of optimization (Bousquet and Bottou, 2008) showing superior runtimes of stochastic gradient descent (SGD) methods compared to their exact counterparts that process the entire data set for every iteration. Notably, although only a small subset of cells are considered for each iteration, each of our gradient updates to *Θ* affects all cells in the data (due to its generalizability). In contrast, applying a similar mini-batch SGD procedure to t-SNE results in an ineffective method, as only the positions of the cells in a given mini-batch are updated while the remaining cells are fixed. This reduces to a coordinate descent-like procedure, which we found to be very slow in terms of learning speed, likely due to the tight coupling of parameters being optimized. Although net-SNE shares the model parameters across all cells and thus is less prone to this issue, we did notice difficulties in optimization with mini-batches that are too small. We found setting |ℬ| to be around 10% of the data set to be a reasonable compromise that leads to good performance.

In addition, sampling strategy for ℬ has a considerable effect on the quality of approximation for the gradients. Specifically, given a sparse approximation of ***P***, the first summation in our equation for *δ*KL(***P*** ∥ ***Q***)/*δ**y***_*i*_ has only a few nonzero summands. If ℬ is uniformly sampled, then only a few indices will contribute to the sum, leading to high variance in the estimate. We address this problem by introducing additional structure into ℬ. In particular, we first sample a smaller set of seed cells 𝒮 ⊂ 1, …, *n* uniformly at random and then sample a fixed number of cells from each of their “neighbors” (where *p*_*ij*_> 0).

After these local samples are used to facilitate the approximation of t-SNE gradients for the seed cells, these gradients are backpropagated through the neural network to update the embedding parameters of net-SNE. Given that our estimated normalization factor 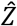 appears in the denominator of the approximate t-SNE gradient, our gradient estimate based on 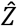 is thus biased. To control the amount of error introduced, we impose a minimum threshold (10%) on the fraction of total samples to be used to approximate 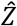. If the set of seed cells is too small, additional samples are drawn to ensure 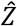 is of sufficient quality.

### Training net-SNE with Reference Visualization

Even with our stochastic optimization techniques, training a neural network to optimize the t-SNE objective is a challenging task, especially for large-scale data sets with complex patterns. We thus introduce another technique, where a pre-trained t-SNE embedding is used to provide more direct feedback to the neural network instead of relying on the highly complex t-SNE objective. More precisely, mean-squared error between the net-SNE embedding and an existing t-SNE map is used as the loss function to optimize the network in order to obtain a good initial solution, which can be further optimized if needed. We found this approach to be very fast and effective. Although this does require t-SNE to be performed before applying net-SNE, we emphasize that the initial t-SNE map need not be fully optimized, as further SGD iterations in net-SNE can fine tune the solution. This approach also suggests the possibility of a more sophisticated one that trains t-SNE and net-SNE *in parallel*, where the t-SNE updates for optimizing the t-SNE objective and the net-SNE updates for closing the gap between the t-SNE and net-SNE outputs are interleaved. Notably, this framework can be viewed as a type of relaxation of our original formulation of net-SNE, in which auxiliary variables are introduced to decouple the two components of the model (i.e., t-SNE optimization and neural network training). Even though mega-scale single cell data sets do not allow us to quickly train t-SNE on the full data set to obtain the initial t-SNE map, we can still use this approach to more effectively train net-SNE on a smaller subset and use the trained embedding to visualize the entire data set via net-SNE’s generalization. This is indeed the approach we take for visualizing the Brain1m data set.

Note that all of our data sets other than PBMC68k and Brain1m are small enough that we were able to train net-SNE *exactly* without stochastic optimization and the reference visualization. On the other hand, our results on PBMC68k and the 100k subset of Brain1m are obtained by training net-SNE to match a t-SNE reference using mini-batch SGD with a batch size of 10%. We did not further fine tune net-SNE after training on the reference visualization, as the resulting visualization quality was sufficiently high. After generalizing the 100k cell visualization to the full Brain1m data set, further training of net-SNE was performed *without* a t-SNE map (since t-SNE becomes impractical at this scale), but we did use our stochastic optimization with batch size of 10%. Although our generalization based on a smaller subset is the primary factor in achieving fast runtime for the Brain1m visualization, our stochastic optimization techniques also lead to significant runtime reductions. For instance, with a batch size of 10%, the average runtime of each iteration of net-SNE is reduced by around a factor of 10 as a result of our techniques.

### Benchmark Data Sets

For our main experiments, we used 13 published scRNA-seq data sets of varying sizes with known cluster labels for the cells, which allowed us to directly assess the quality of the visualization produced by net-SNE. The list of data sets, sorted in increasing order of size: (Biase et al., 2014) (*n* = 49, *k* = 3), (Treutlein et al., 2014) (*n* = 80, *k* = 5), (Goolam et al., 2016) (*n* = 124, *k* = 5), (Ting et al., 2014) (*n* = 149, *k* = *7*), (Buettner et al., 2015) (*n* = 182, *k* = 3), (Deng et al., 2014) (*n* = 268, *k* = 10), (Pollen et al., 2014) (*n* = 301, *k* = 11), (Patel et al., 2014) (*n* = 430, *k* = 5), (Usoskin et al., 2015) (*n* = *622*, *k* = 4), (Kolodziejczyk et al., 2015) (*n* = 704, *k* = 3), (Klein et al., 2015) (*n* = *2*,717, *k* = 4), (Zeisel et al., 2015) (*n* = 3,005, *k* = 9), and PBMC68k (Zheng et al., 2017) (*n* = 68,560, *k* = 10). Note that *n* denotes the number of cells and *k* denotes the number of clusters defined by the original publications.

We additionally used a mega-scale data set, Brain1m (10x Genomics, 2017) (*n* = 1,283,543), to assess the generalizability and scalability of net-SNE. However, this data set does not have any known labels, and thus we resorted to using the objective score optimized by both net-SNE and t-SNE as the quality metric for comparison.

Our benchmark data sets measured gene expression via a number of different metrics, including read, fragment, transcript, or unique molecule (UMI) counts that are either normalized or unnormalized for gene length (Table S1). We kept the original metric chosen by the original publication with the goal of demonstrating the performance of net-SNE in a variety of settings. Unless already performed by the original publication, we applied the standard log (1 + *x*) transformation to each element *x* of the cell-gene expression matrix before visualizing the data.

### Data and Software Availability

A C++ implementation of net-SNE is available at: http://netsne.csail.mit.edu and https://github.com/hhcho/netsne.

## References

10x Genomics, 2017. Transcriptional Profiling of 1.3 Million Brain Cells with the Chromium Single Cell 3’ Solution.

Amir, E.-A.D., Davis, K.L., Tadmor, M.D., Simonds, E.F., Levine, J.H., Bendall, S.C., Shenfeld, D.K., Krishnaswamy, S., Nolan, G.P., Pe’er, D., 2013. viSNE enables visualization of high dimensional single-cell data and reveals phenotypic heterogeneity of leukemia. Nature biotechnology 31, 545.

Anchang, B., Hart, T.D.P., Bendall, S.C., Qiu, P., Bjornson, Z., Linderman, M., Nolan, G.P., Plevritis, S.K., 2016. Visualization and cellular hierarchy inference of single-cell data using SPADE. Nature Protocols 11, 1264–1279. doi:10.1038/nprot.2016.066

Biase, F.H., Cao, X., Zhong, S., 2014. Cell fate inclination within 2-cell and 4-cell mouse embryos revealed by single-cell RNA sequencing. Genome research 24, 1787–1796.

Bousquet, O., Bottou, L., 2008. The tradeoffs of large scale learning, in:. Presented at the Advances in neural information processing systems, pp. 161–168.

Buettner, F., Natarajan, K.N., Casale, F.P., Proserpio, V., Scialdone, A., Theis, F.J., Teichmann, S.A., Marioni, J.C., Stegle, O., 2015. Computational analysis of cell-to-cell heterogeneity in single-cell RNA-sequencing data reveals hidden subpopulations of cells. Nature biotechnology 33, 155–160.

Deng, Q., Ramsköld, D., Reinius, B., Sandberg, R., 2014. Single-cell RNA-seq reveals dynamic, random monoallelic gene expression in mammalian cells. Science 343, 193–196.

Dzwinel, W., Wcisło, R., 2015. Very Fast Interactive Visualization of Large Sets of High-dimensional Data. Procedia Computer Science 51, 572–581. doi:https://doi.org/10.1016/j.procs.2015.05.325

Gawad, C., Koh, W., Quake, S.R., 2016. Single-cell genome sequencing: current state of the science. Nature Reviews Genetics 17, 175–188.

Goolam, M., Scialdone, A., Graham, S.J., Macaulay, I.C., Jedrusik, A., Hupalowska, A., Voet, T., Marioni, J.C., Zernicka-Goetz, M., 2016. Heterogeneity in Oct4 and Sox2 targets biases cell fate in 4-cell mouse embryos. Cell 165, 61–74.

Grün, D., Lyubimova, A., Kester, L., Wiebrands, K., Basak, O., Sasaki, N., Clevers, H., van Oudenaarden, A., 2015. Single-cell messenger RNA sequencing reveals rare intestinal cell types. Nature 525, 251–255.

Haghverdi, L., Lun, A.T.L., Morgan, M.D., Marioni, J.C., 2017. Correcting batch effects in single-cell RNA sequencing data by matching mutual nearest neighbours. bioRxiv. doi:10.1101/165118

Hartigan, J.A., Wong, M.A., 1979. Algorithm AS 136: A k-means clustering algorithm. Journal of the Royal Statistical Society. Series C (Applied Statistics) 28, 100–108.

Hutchison, L.A.D., Berger, B., Kohane, I., 2017. C. elegans exhibits coordinated oscillation in gene expression during development. bioRxiv. doi:10.1101/114074

Jackson, J.E., 2005. A user’s guide to principal components. John Wiley & Sons.

Jaitin, D.A., Kenigsberg, E., Keren-Shaul, H., Elefant, N., Paul, F., Zaretsky, I., Mildner, A., Cohen, N., Jung, S., Tanay, A., others, 2014. Massively parallel single-cell RNA-seq for marker-free decomposition of tissues into cell types. Science 343, 776–779.

Kiselev, V.Y., Kirschner, K., Schaub, M.T., Andrews, T., Yiu, A., Chandra, T., Natarajan, K.N., Reik, W., Barahona, M., Green, A.R., others, 2017. SC3: consensus clustering of single-cell RNA-seq data. Nature Methods.

Klein, A.M., Mazutis, L., Akartuna, I., Tallapragada, N., Veres, A., Li, V., Peshkin, L., Weitz, D.A., Kirschner, M.W., 2015. Droplet barcoding for single-cell transcriptomics applied to embryonic stem cells. Cell 161, 1187–1201.

Kolodziejczyk, A.A., Kim, J.K., Tsang, J.C., Ilicic, T., Henriksson, J., Natarajan, K.N., Tuck, A.C., Gao, X., Bühler, M., Liu, P., others, 2015. Single cell RNA-sequencing of pluripotent states unlocks modular transcriptional variation. Cell Stem Cell 17, 471–485.

LeCun, Y., Bengio, Y., Hinton, G., 2015. Deep learning. Nature 521, 436–444.

Loh, P.-R., Baym, M., Berger, B., 2012. Compressive genomics. Nature biotechnology 30, 627–630.

Maaten, L.V.D., Hinton, G., 2008. Visualizing data using t-SNE. Journal of Machine Learning Research 9, 2579–2605.

Moon, K.R., van Dijk, D., Wang, Z., Burkhardt, D., Chen, W., van den Elzen, A., Hirn, M.J., Coifman, R.R., Ivanova, N.B., Wolf, G., Krishnaswamy, S., 2017. Visualizing Transitions and Structure for High Dimensional Data Exploration. bioRxiv. doi:10.1101/120378

Palmer, N.P., Schmid, P.R., Berger, B., Kohane, I.S., 2012. A gene expression profile of stem cell pluripotentiality and differentiation is conserved across diverse solid and hematopoietic cancers. Genome biology 13, R71.

Patel, A.P., Tirosh, I., Trombetta, J.J., Shalek, A.K., Gillespie, S.M., Wakimoto, H., Cahill, D.P., Nahed, B.V., Curry, W.T., Martuza, R.L., others, 2014. Single-cell RNA-seq highlights intratumoral heterogeneity in primary glioblastoma. Science 344, 1396–1401.

Pierson, E., Yau, C., 2015. ZIFA: Dimensionality reduction for zero-inflated single-cell gene expression analysis. Genome biology 16, 241.

Pollen, A.A., Nowakowski, T.J., Shuga, J., Wang, X., Leyrat, A.A., Lui, J.H., Li, N., Szpankowski, L., Fowler, B., Chen, P., others, 2014. Low-coverage single-cell mRNA sequencing reveals cellular heterogeneity and activated signaling pathways in developing cerebral cortex. Nature biotechnology 32, 1053–1058.

Qiu, X., Mao, Q., Tang, Y., Wang, L., Chawla, R., Pliner, H.A., Trapnell, C., 2017. Reversed graph embedding resolves complex single-cell trajectories. Nature Methods 14, 979–982.

Rand, W.M., 1971. Objective criteria for the evaluation of clustering methods. Journal of the American Statistical association 66, 846–850.

Regev, A., Teichmann, S., Lander, E.S., Amit, I., Benoist, C., Birney, E., Bodenmiller, B., Campbell, P., Carninci, P., Clatworthy, M., others, 2017. The Human Cell Atlas. bioRxiv 121202.

Samet, H., 1984. The quadtree and related hierarchical data structures. ACM Computing Surveys (CSUR) 16, 187–260.

Simmons, S., Peng, J., Bienkowska, J., Berger, B., 2015. Discovering What Dimensionality Reduction Really Tells Us About RNA-Seq Data. Journal of Computational Biology 22, 715–728.

Stubbington, M.J., Rozenblatt-Rosen, O., Regev, A., Teichmann, S.A., 2017. Single-cell transcriptomics to explore the immune system in health and disease. Science 358, 58–63.

Svensson, V., Vento-Tormo, R., Teichmann, S.A., 2017. Exponential scaling of single-cell RNA-seq in the last decade. arXiv preprint arXiv:1704.013795.

Tang, J., Liu, J., Zhang, M., Mei, Q., 2016. Visualizing Large-scale and High-dimensional Data. Proceedings of the 25th International Conference on World Wide Web 287–297. doi:10.1145/2872427.2883041

Ting, D.T., Wittner, B.S., Ligorio, M., Jordan, N.V., Shah, A.M., Miyamoto, D.T., Aceto, N., Bersani, F., Brannigan, B.W., Xega, K., others, 2014. Single-cell RNA sequencing identifies extracellular matrix gene expression by pancreatic circulating tumor cells. Cell reports 8, 1905–1918.

Treutlein, B., Brownfield, D.G., Wu, A.R., Neff, N.F., Mantalas, G.L., Espinoza, F.H., Desai, T.J., Krasnow, M.A., Quake, S.R., 2014. Reconstructing lineage hierarchies of the distal lung epithelium using single-cell RNA-seq. Nature 509, 371–375.

Tung, P.-Y., Blischak, J.D., Hsiao, C.J., Knowles, D.A., Burnett, J.E., Pritchard, J.K., Gilad, Y., 2017. Batch effects and the effective design of single-cell gene expression studies. Scientific reports 7, 39921.

Usoskin, D., Furlan, A., Islam, S., Abdo, H., Lönnerberg, P., Lou, D., Hjerling-Leffler, J., Haeggström, J., Kharchenko, O., Kharchenko, P.V., others, 2015. Unbiased classification of sensory neuron types by large-scale single-cell RNA sequencing. Nature neuroscience 18, 145–153.

Van Der Maaten, L., 2014. Accelerating t-SNE Using Tree-based Algorithms. Journal of Machine Learning Research 15, 3221–3245.

Van Der Maaten, L., 2009. Learning a parametric embedding by preserving local structure. RBM 500, 26.

Wang, B., Zhu, J., Pierson, E., Ramazzotti, D., Batzoglou, S., 2017. Visualization and analysis of single-cell RNA-seq data by kernel-based similarity learning. Nature Methods 14, 414–416.

Wang, Y., Waters, J., Leung, M.L., Unruh, A., Roh, W., Shi, X., Chen, K., Scheet, P., Vattathil, S., Liang, H., others, 2014. Clonal evolution in breast cancer revealed by single nucleus genome sequencing. Nature 512, 155–160.

Yoon, H.S., Price, D.C., Stepanauskas, R., Rajah, V.D., Sieracki, M.E., Wilson, W.H., Yang, E.C., Duffy, S., Bhattacharya, D., 2011. Single-cell genomics reveals organismal interactions in uncultivated marine protists. Science 332, 714–717.

Yu, Y.W., Daniels, N.M., Danko, D.C., Berger, B., 2015. Entropy-scaling search of massive biological data. Cell Systems 1, 130–140.

Zeisel, A., Muñoz-Manchado, A.B., Codeluppi, S., Lönnerberg, P., La Manno, G., Juréus, A., Marques, S., Munguba, H., He, L., Betsholtz, C., others, 2015. Cell types in the mouse cortex and hippocampus revealed by single-cell RNA-seq. Science 347, 1138–1142.

Zheng, G.X., Terry, J.M., Belgrader, P., Ryvkin, P., Bent, Z.W., Wilson, R., Ziraldo, S.B., Wheeler, T.D., McDermott, G.P., Zhu, J., others, 2017. Massively parallel digital transcriptional profiling of single cells. Nature communications 8, 14049.

