## Supplementary Materials for "Neural Data Visualization for Scalable and Generalizable Single Cell Analysis"

| **Data set** | **N** | **K** | **Count type** | **Cell type** | **Publication** |
|---|---|---|---|---|---|
| Biase | 49 | 3 | FPKM | Mouse embryo | 2014 |
| Treutlein | 80 | 5 | FPKM | Mouse lung | 2014 |
| Goolam | 124 | 5 | CPM | Mouse embryo | 2016 |
| Ting | 149 | 7 | CPM | Mouse pancreatic tumor | 2014 |
| Buettner | 182 | 3 | FPKM | Mouse embryonic stem cells | 2015 |
| Deng | 268 | 10 | RPKM | Mouse embryo | 2014 |
| Pollen | 301 | 11 | TPM | Human brain | 2014 |
| Patel | 430 | 5 | TPM | Human brain tumor | 2014 |
| Usoskin | 622 | 4 | RPM | Mouse brain | 2014 |
| Kolodziejczyk | 704 | 3 | CPM | Mouse embryonic stem cells | 2015 |
| Klein | 2,717 | 4 | UMI | Mouse embryonic stem cells | 2015 |
| Zeisel | 3,005 | 9 | UMI | Mouse brain | 2015 |
| PBMC68k | 68,560 | 10 | UMI | Human blood | 2017 |
| Brain1m | 1,283,543 | - | UMI | Mouse brain | 2017 |

**Table S1.** Related to Figure 2. List of publicly available single cell RNA-seq data sets we used to evaluate net-SNE, in increasing order of size. N denotes the number of cells, and K denotes the number of clusters used in the annotations provided by the original publications. Count type refers to the metric used to represent the expression level of each gene in each data set. Our benchmark data sets cover a wide range of cell types and analysis settings. FPKM: fragments per kilobase of transcript per million mapped reads. CPM: counts per million mapped reads. RPKM: reads per kilobase of transcript per million mapped reads. TPM: transcripts per million mapped reads. RPM: reads per million mapped reads. UMI: unique molecular identifier counts. Reference for each data set is provided in Methods.

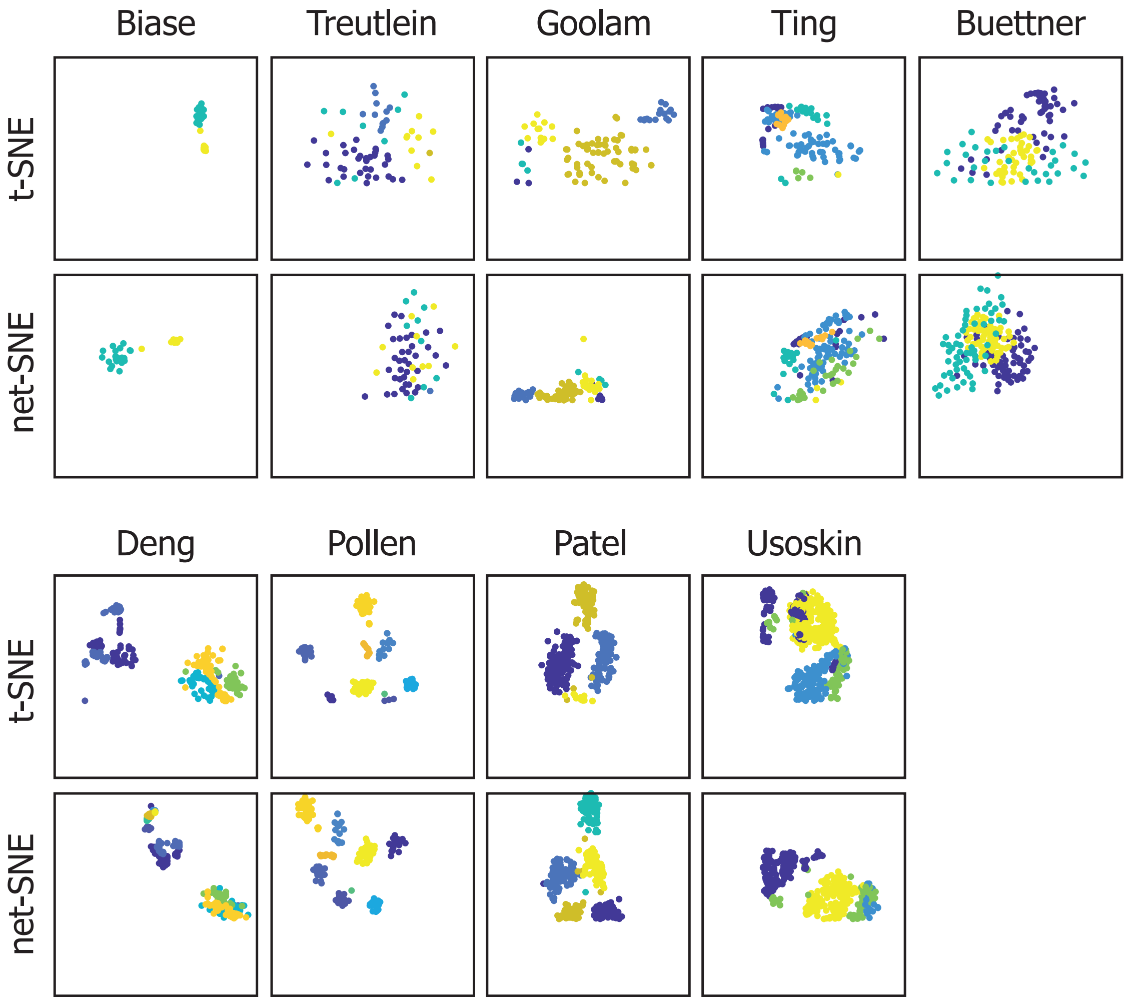


**Figure S1.** Related to Figure 2. net-SNE recapitulates t-SNE mapping on 13 benchmark data sets with known subtypes. The remaining 4 data sets are shown in Figure 2. Each point represents a cell, and the colors correspond to cluster assignments provided by the original publications. The references for our benchmark data sets are provided in Methods.

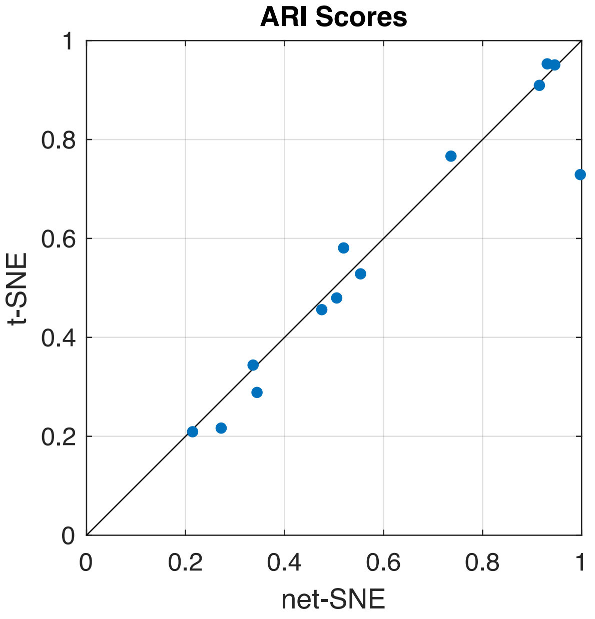


**Figure S2.** Related to Figure 2. We quantified the quality of net-SNE visualization on 13 benchmark data sets with known cluster labels by the adjusted Rand index (ARI) between the reference clustering and the output of agglomerative clustering (based on Euclidean distance) of the cells in the 2D visualization. Each dot in the scatter plot represents a distinct benchmark data set. Overall, net-SNE achieves clustering accuracies that closely match those of t-SNE, with the exception of the Kolodziejczyk data set, on which net-SNE outperforms t-SNE. Alternative results based on k-means clustering are provided in Figure 2b.

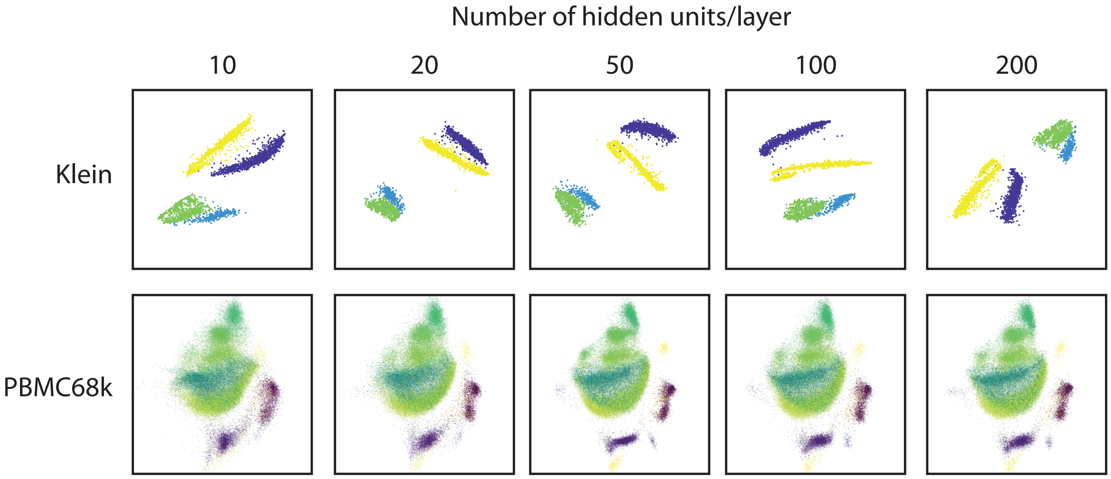


**Figure S3.** Related to Figure 2. net-SNE visualization is robust to a reasonably wide range of neural network sizes for modeling the embedding function. Note that all our experiments are based on a relatively shallow neural network with two layers. As shown, varying the number of hidden units in each layer from 10 to 200 did not significantly affect the visualization quality, although a moderate amount of blur can be seen in the visualization of PBMC68k data set using 10 units/layer, which can be attributed to limited model capacity. The fact that PBMC68k visualizations maintain the same global organization unlike the Klein data set is due to the fact that we used the same t-SNE map to guide and accelerate net-SNE optimization for PBMC68k (Methods). In contrast, net-SNE trains without guidance on the Klein data set, thereby resulting in different (albeit mostly symmetric) configurations of the clusters that depend on the initializations.

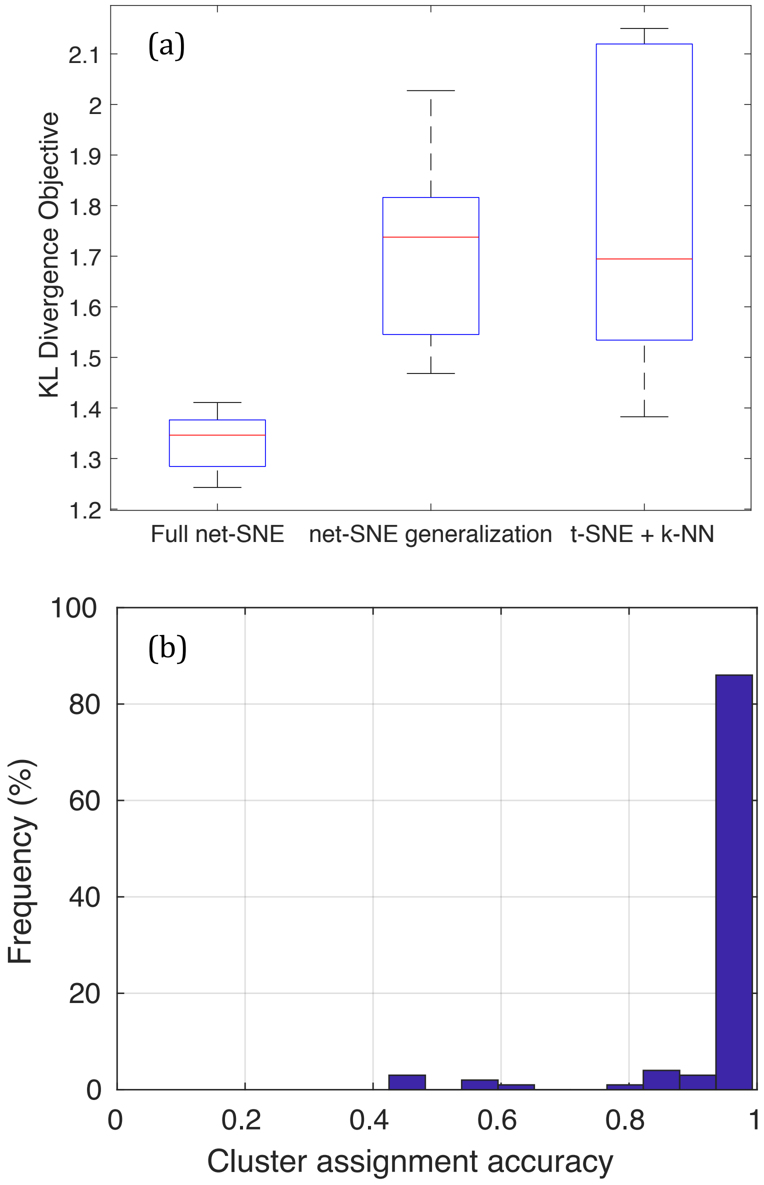


**Figure S4.** Related to Figure 3. (a) On the Klein data set, we performed a cross validation experiment where each time cells that belong to one of the four known clusters are entirely held out from the initial visualization by net-SNE or t-SNE and later added to the image, using the trained net-SNE embedding for net-SNE or a k-nearest neighbor approach for t-SNE (i.e., positioning each new cell at the average location of its five closest cells in the gene expression space). We quantified the quality of the resulting joint visualization by the Kullback-Leibler (KL) divergence score on the full data set that both net-SNE and t-SNE minimize. Although the difference between net-SNE's generalization (center) and the k-NN extension of t-SNE (right) was statistically inconclusive (Wilcoxon rank sum p-value > 0.1), clearly better visual quality of net-SNE's generalization (shown in Figure 3a) is consistent with the increased rate of poor objective values (around 2) for t-SNE with k-NN, which is a result of the unseen cluster being spread between two other clusters, due to the limitations of k-NN. Interestingly, when the new cluster is completely enclosed in another cluster (which renders it invisible), the resulting KL objective was reasonably good, thereby neutralizing the overall performance of t-SNE with k-NN. Training net-SNE on the full data from scratch (left) achieved better objective scores than both generalization approaches, as expected. However, we note that such a joint visualization becomes impractical when the scale of data sets reach millions of cells. Boxes summarize the distribution of results from 20 trials. Red mark indicates the median. Box edges indicate the lower and upper quartiles. Whiskers extend to the most extreme values in the distribution. (b) To evaluate the accuracy of net-SNE generalization in a setting where cells that belong to the same subtype as the new cell are already represented in the initial data set, we performed a two-fold cross validation analysis. For each new cell, we used the net-SNE embedding to find its position and predicted its cluster based on a majority vote of nearest neighbors in the visualization. Histogram of the fraction of new cells that were assigned to the correct cluster is shown, which summarizes the data from 100 trials. With high frequency (>80%), net-SNE correctly positions almost all of the new cells to where their corresponding subtypes are represented.
